# The Centrosomal Swiss Army Knife: A combined in silico and in vivo approach to the structure-function annotation of SPD-2 provides mechanistic insight into its functional diversity

**DOI:** 10.1101/2021.04.22.441031

**Authors:** Mikaela Murph, Shaneen Singh, Mara Schvarzstein

## Abstract

Centrosomes are organelles that function as hubs of microtubule nucleation and organization, with key roles in organelle positioning, asymmetric cell division, and ciliogenesis. Aberrant centrosome structure or function is linked to neurodegenerative diseases, developmental abnormalities, ciliopathies, and tumor development. A major regulator of centrosome biogenesis and function in *C. elegans* is the highly conserved protein Spindle-defective protein 2 (SPD-2), a homolog of the human CEP-192 protein. CeSPD-2 is required for centrosome maturation, centriole duplication, spindle assembly and cell polarity establishment. Despite its importance, the specific molecular mechanism of CeSPD-2 function is poorly understood. To address this gap in knowledge, we combined computational analysis with cell biology approaches to uncover structure-function relationships of CeSPD-2 that may shed mechanistic light on its function. Domain prediction analysis corroborated and refined previously identified coiled-coils and ASH (Aspm-SPD-2 Hydin) domains and identified new domains and motifs: an additional coiled-coil, a GEF domain, an Ig-like domain, and a PDZ-like domain. Our findings suggest that ASH domain belongs to the same superfold as PapD chaperone domains and Major Sperm Protein (MSP) domains within the larger Immunoglobulin superfamily. We have identified a large novel basic region in the CeSPD-2 ASH domain that harbors most of the predicted protein and nucleic acid contact residues in the ASH domain. *In vivo*, ASH::GFP localized to centrosomes and centrosome associated microtubules, and forms aggregates in the cytosol when overexpressed. This study lays the groundwork for designing rational hypothesis-based experiments for future analyses to further elaborate the mechanisms of CeSPD-2 function *in vivo*.

## Introduction

Centrosomes are organelles that play key roles in the positioning of other organelles and function as hubs for microtubule nucleation and organization. They are composed of a pair of microtubular centrioles that are immersed in a protein matrix known as the pericentriolar material (PCM). Despite the codependence between the centrioles and the PCM for many of the centrosome functions, they have different roles and largely associate with a distinct set of proteins^1^. An exception is the *C. elegans* spindle-defective protein 2 (SPD-2) protein that is crucial for centriole duplication and centrosome maturation and localizes either to the centrosome or exclusively to the centrioles ^2,3^. This dynamic localization is likely relevant to CeSPD-2’s dual function regulating microtubule nucleation and centriole duplication ^2^.

In mitosis the centrosome dynamics and cell cycle progression are coordinated to ensure a single centrosome duplication per cell cycle^4^. At fertilization in the *C. elegans* the sperm contributes a pair of centrioles and the oocyte contributes the PCM, and together they form the first centrosome pair of the one-cell embryo. Accumulation of maternally provided PCM in the zygote is promoted by the sperm provided centrioles. PCM accumulates microtubule nucleating proteins such as the γ-tubulin microtubule anchoring protein and the microtubule polymerase ZYG-9/MAP215. PCM assembly in the *C elegans* embryo occurs in steps during the cell cycle^1^. CeSPD-2 is required in all these steps. At fertilization, SPD-2 localizes to the centriole pair and in a shell around the sperm DNA^5^. Soon after the centriole pair separate, PCM starts to accumulate around them. PCM assembly requires SAS-7, which is also required for the assembly of the daughter centrioles^6^. The coiled-coil protein PCM deficient-1 (PCMD-1) is required for the recruitment of SPD-5^7^. PCMD-1, SPD-5, SPD-2 and γ-tubulin are the earliest detectable PCM proteins accumulating around the disengaged pair of centrioles provided by the sperm. This occurs as anteroposterior polarity is being established, a process that depends on the maturing centrosome and possibly on SPD-2^8^. Once the embryo enters mitosis, the PCM volume and γ-tubulin content increases several-fold^9,10^. SPD-2 accumulation at the centrosome is partially dependent on AIR-1 and the cytoplasmic dynein *dhc-1*^2^. Together SPD-2 and SPD-5 promote PCM assembly and centrosome maturation^2^. SPD-2 recruits polo-like kinase-1 (PLK-1), and Aurora kinase A/AIR-1 to the PCM, which are required for centrosome maturation^11,12^. At metaphase PCM components are restructured. PLK-1 phosphorylates SPD-5^13^ and this initiates polymerization of PCM and the formation of a γ-tubulin ring surrounded by a microtubule ring^14,15^. Spindle microtubules emanate from the interphase between these two concentric rings^16^.

After fertilization, the sperm provided centrioles separate they start to assemble daughter centrioles by sequentially recruiting maternally provided structural and regulatory centriolar components^17–19^, a process that also requires PCM proteins^20^. Six proteins have been shown to be required for centriole assembly; the zygote defective 1 protein (ZYG-1) ^21^, the likely ortholog of Polo like kinase 4 (PLK4) in mammals, and five coiled-coil proteins: SPD-2^2,3^, and Spindle assembly abnormal proteins SAS-4^22^, SAS-5^23^, SAS-6^24^ and SAS-7^6^. SAS-7 is thought to directly bind to and recruit CeSPD-2 to the emerging daughter centrioles on the separated centrioles^6^. The N-terminal acidic region of CeSPD-2 binds to the basic region on ZYG-1/PLK4 kinase, which in turn recruits SAS-5 and together they form the SAS-6 containing cartwheel organization on the daughter centriole^18,25^. SAS-4 localization to the daughter centriole seems to take place concomitantly with SAS-6 and regulates the formation of the microtubule containing outer wall of the centriole^18^. This is followed by the formation of the SAS-7 promoted ninefold symmetric paddlewheel completing the process of the formation of the daughter centrioles^6^.

The mechanisms and protein machinery required for centriole formation in the centriole duplication process are highly conserved, with the minimum core of proteins essential for centriole biogenesis in all organisms studied include orthologs of ZYG-1/PLK4, SAS-4/CPAP, and SAS-6^26^. *C. elegans* SPD-2 and its human ortholog CEP192 are also essential for centriole duplication^3,27,28^, however it is not clear whether DsSPD-2 is essential for centriole duplication in spermatocytes^29,30^.

The myriad functions and intermolecular interactions of SPD-2 during the cell cycle together with a poor understanding of its mechanisms of function makes SPD-2 an excellent target for studying the mechanisms that underlie centrosome biogenesis and inheritance. In this study, we report that despite being predicted to be an intrinsically disordered protein, CeSPD-2 C-terminal half has structured regions with several functional domains including a previously unreported fourth coiled-coil, a GEF domain, an ASH domain, an Ig-like domain, and a PDZ-like domain. The predicted three-dimensional structure of the SPD-2 ASH domain reveals that it is a member of a structural superfold that includes Major Sperm Protein (MSP), PapD, and usher-chaperone domains. Our studies also show that SPD-2 ASH, localizes to centrosomes and centrosome associated microtubules in *C. elegans* embryos. Overall, this study lays the groundwork for designing hypothesis-based experiments for future analyses to probe the suggested mechanisms of SPD-2 function *in vivo*.

## Results

### CeSPD-2 is a Multidomain Protein with an Intrinsically Disordered N-terminus

The predicted secondary structure of *C. elegans* SPD-2 protein sequence (NP_492414.1), suggests that it is 50% unstructured (Figure 1 and 2). The first 460 residues in CeSPD-2 are predicted to be 72% intrinsically disordered (72%), suggesting that this region may not form stable structures independently (Figure 2). Consistent with this the secondary structure of the N-terminal half of SPD-2 is predicted to yield only five to seven α-helices (Figure 1). However, the C-terminal half of the protein is predicted to contain several secondary structure signatures indicating hallmarks of structured domains. CeSPD-2 has been previously reported to have three coiled-coils^2^. Our analysis identified an additional coiled-coil region (368-400) (Figure 2) that was not previously reported. We also predict CC3 and CC4 to be trimeric coiled-coils with a potential for oligomerization (Figures 1 and 2); they lie within a predicted RhoGEF domain motif (258–467) with similarity to the hypothetical Rom1 protein from *Encephalitozoon cuniculi*. CC3 and CC4 appear to be analogous to the catalytic coiled-coils that form the Sec2p GEF domain fold (Figure 2)^31^. The structured C-terminal region of CeSPD-2 houses an Aspm-SPD-2-Hydin (ASH) domain (475 – 564), followed by an Immunoglobulin (Ig)-like domain (593-660) made up of six stable antiparallel *β*-sheets in a sandwich conformation structurally similar to the S-type Immunoglobulin fold, and a putative PDZ-like domain (695 - 820) containing a carboxylate-binding loop region [R/K-X-X-X-Φ-G-Φ] characteristic of PDZ/PDZ-like domains^32,33^ (R743 - F749) of the domain (Figures 1 and 2).

**Figure 1.**
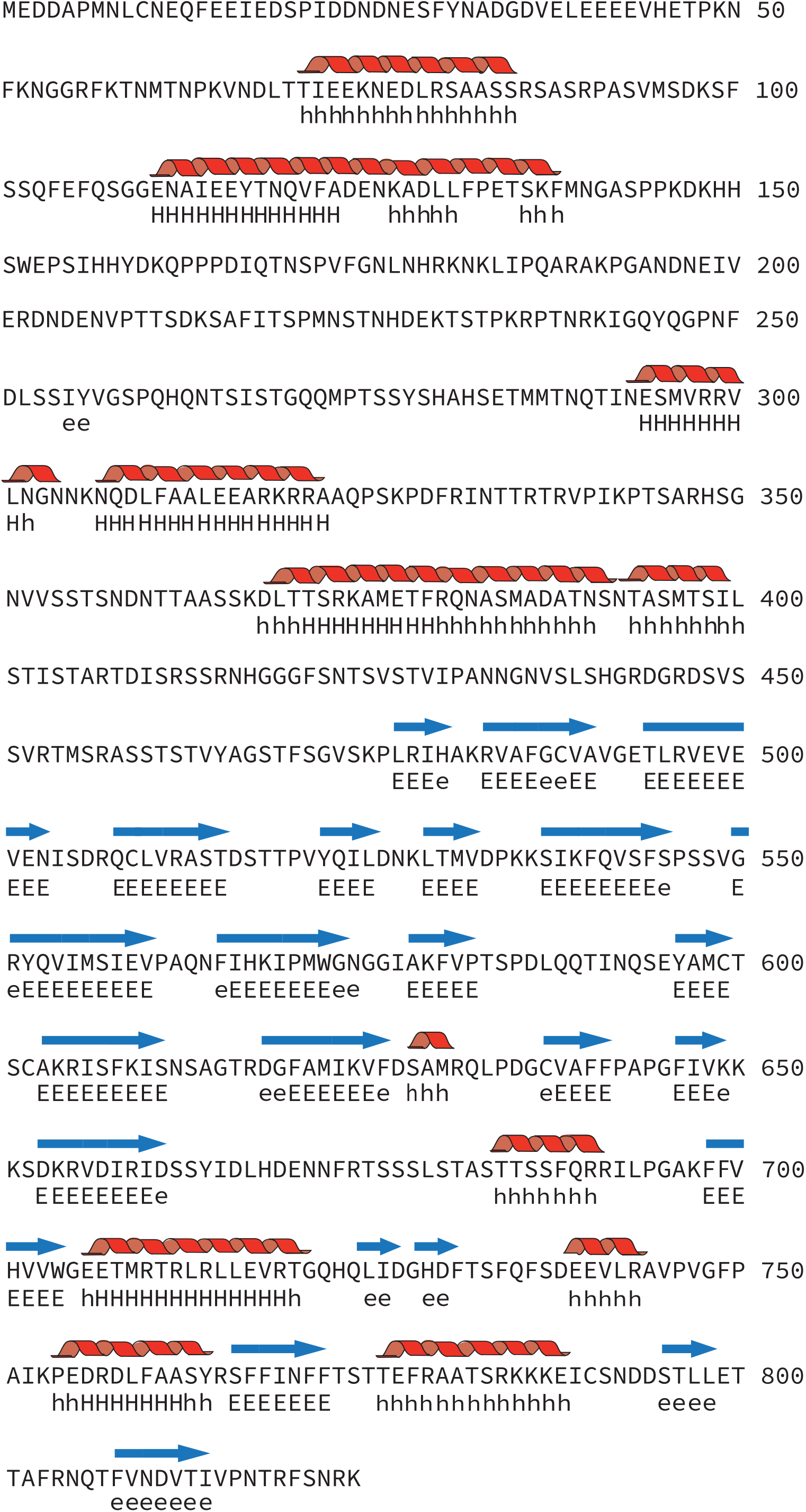
Secondary structure prediction of CeSPD-2. The predicted secondary structure of SPD-2 based on a consensus of multiple prediction programs illustrated onto its the primary amino acid sequence. CeSPD-2 is predicted to include 11 α-helices (represented by red ribbons) and 21 β-strands (represented by blue arrows). The α-helices are also represented by an H or an h, and β-strands are represented by an E or an e. Lowercase letters (h and e) refers to residues that are predicted to form structures that meet at least a probability threshold of 0.5, and uppercase letters (H and E) refers to residues that are equal or exceed the probability threshold of 0.6.

**Figure 2.**
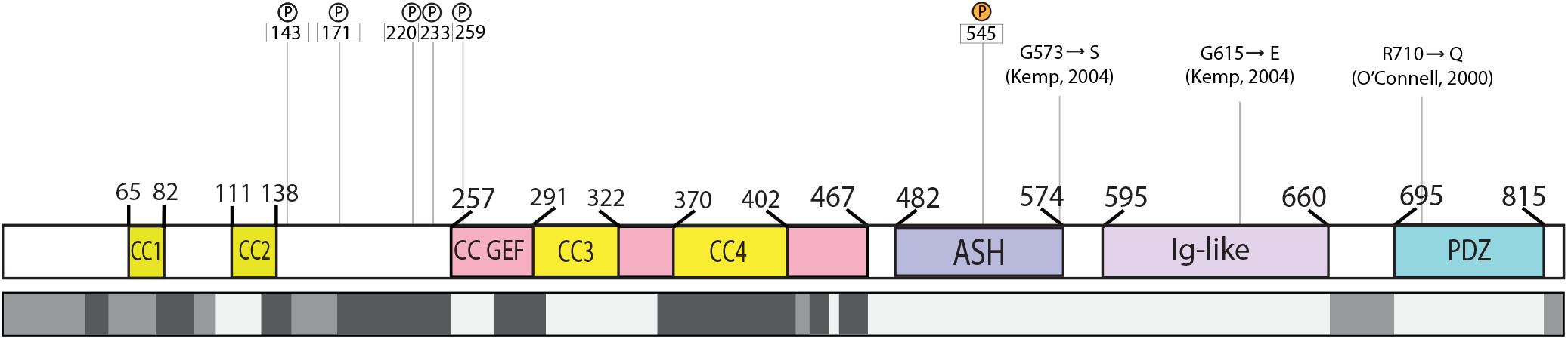
Predicted domain architecture of CeSPD-2. To scale illustration of the structural domains of CeSPD-2 inferred from literature^40^, and sequence analysis. There are four predicted coiled-coil regions (labelled CC1 – CC4), two of which (CC3 and CC4) are part of the predicted GTPase guanine nucleotide exchange factor (GEF) domain. Also illustrated are the locations of an ASPM-SPD2-Hydin (ASH) domain, an Immunoglobulin-like/Major Sperm Protein (Ig-like/MSP) domain and a PSD95/Dlg-1/zo-1 (PDZ) domain. Amino acid positions in the protein are provided at the limits of each domain for cross-reference. Regions predicted to be disordered are shown underneath in a grayscale bar, where the degree of disorder is represented by light gray (low, < 0.5), medium gray (0.5 - 0.6), and dark gray (high, ≥0.6). Selected known alleles (S545E, G573S, G615E and R710Q) and CDK are also indicated.

### ASH domains show low conservation of primary structure, but have a highly conserved underlying secondary structure

A PSI BLAST search (E-value threshold = 0.005) using the CeSPD-2 ASH domain identified only three significant sequence matches besides the predicted *Caenorhabditis* species homologs of SPD-2 from *C. brigssae*, *C. remanei*, and the hookworm parasite *Necator americanus*. The third iteration identified 11 additional hypothetical or predicted protein sequences at or above 0.05 threshold (Table 1). Given the lack of identification of ASH domains from proteins with known function from the PSI BLAST search, we decided to compare the CeSPD-2 ASH domain sequence to the known ASH domains from characterized proteins known to have roles in ciliary, flagellar, and centrosomal functions including the centrosomal proteins *Homo sapiens* and *Xenopus laevis* Centrosomal Protein 192 (Cep192), *Drosophila melanogaster* Abnormal Spindle Protein (ASP), and human Abnormal Spindle-like Microcephaly-associated Protein (ASPM)^34–38^, the ciliary protein Hydin (*Homo sapiens* and *Mus musculus*), and the *Homo sapiens* Golgi endocytic trafficking protein Inositol polyphosphate 5-phosphatase (OCRL-1)^39^. Alignment of these ASH domains shows that they share little sequence similarity (Figure 3A). CeSPD-2 ASH is most similar (33.33%) to the non-orthologous ASH domain in ASPM from humans (Figure 3B) and CeSPD-2 ASH and its orthologs CEP192 in humans and *Xenopus* only share 26.88% and 32.39% similarity respectively. The only strongly conserved amino acid is an asparagine (N503 in SPD-2) that has been previously defined as a signature feature of ASH domains^40^ (Figure 3A). Other highly conserved residues include an N-terminal phenylalanine (F486 in CeSPD-2 but absent in XeCEP192), and a proline (P534 in SPD-2, absent in ASP and HsCEP192). Comparison of the OCRL-1 ASH domain (3QIS) ^41^ and the predicted ASH domain of CeSPD-2 showed that they have 7 *β*-strands located at similar positions within the sequence (Figure 3A).

**Table 1.**
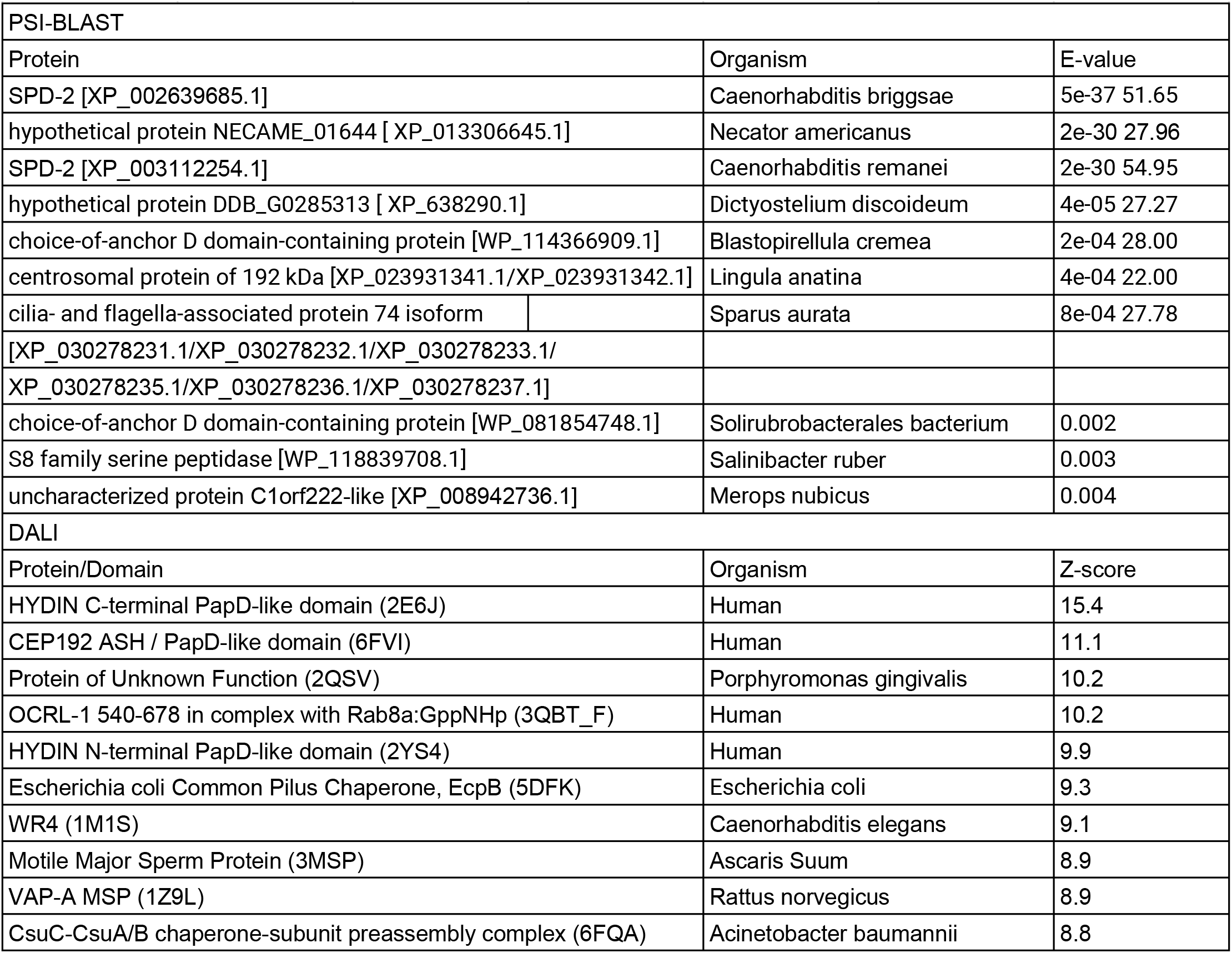
CeSPD-2 ASH Sequence and Structural Homology. A) PSI-BLAST search (third iteration) against the RefSeq database using the *C. elegans* SPD-2 ASH domain sequence. B) The top ten structural homologues of the modeled CeSPD-2 ASH domain, identified by DALI.

**Figure 3.**
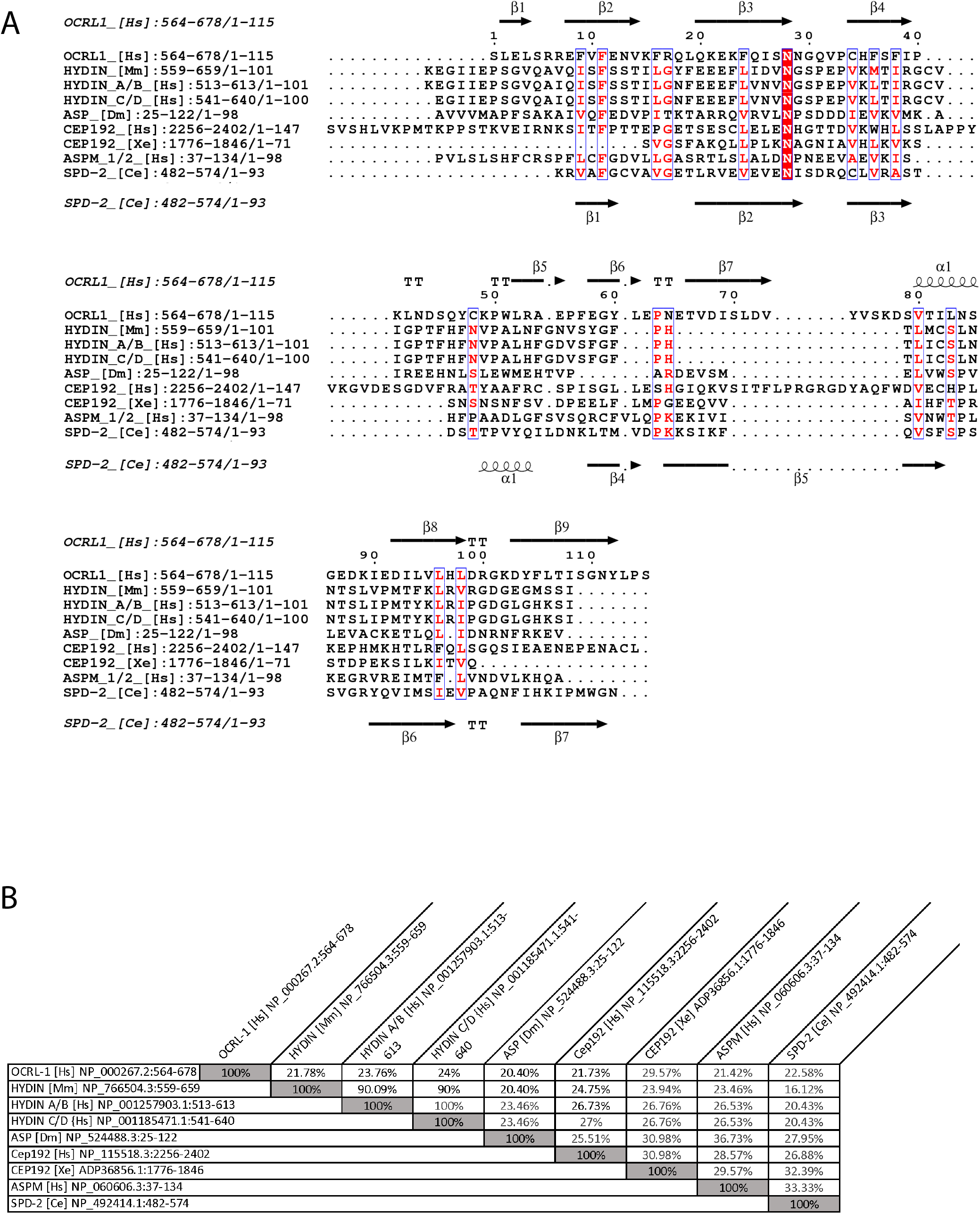
Comparison of sequences of ASH domains from centrosomal proteins. A) Multiple sequence alignment of the ASH domains for centrosomal proteins from *C. elegans*, *Xenopus*, *Drosophila*, *Mus musculus* and *Homo sapiens* using the Muscle algorithm and annotated with ESPript^122^. The sequences shown include *Caenorhabditis elegans* [Ce] spindle defective protein-2/SPD-2 (NP_492414.1:482-574); Human [Hs] hydrocephalus-inducing protein/HYDIN isoforms a/b and c/d (NP_001257903.1:513-613, NP_001185471.1:541-640), abnormal spindle-like microcephaly-associated protein/ASPM isoform 1/2 (NP_060606.3:37-134), inositol polyphosphate 5-phosphatase/OCRL-1 isoform a (NP_000267.2:564-678) and centrosomal protein 192kDa/CEP192 (NP_115518.3:2256-2402); *Xenopus laevis* [Xe] centrosomal protein/CEP192 (ADP36856.1:1776-1846); *Mus musculus* [Mm] hydrocephalus inducing protein/HYDIN (NP_766504.3:559-659); *Drosophila melanogaster* [Dm] abnormal spindle protein (NP_524488.3:25-122). Amino acids with 70% equivalency are labeled in red and boxed in blue. Highly conserved residues are highlighted red. Also shown is the location of secondary structures in the context of the primary sequence for CeSPD-2 and the OCRL-1 ASH domains. In the secondary structure diagrams for CeSPD-2 and OCRL-1, β-sheets are shown as arrows and the α-helices are shown as corkscrews. B) Sequence Similarity of ASH Domains depicted in a similarity matrix comparing the nine observed ASH domains. The matrix was constructed using SIAS.

### The modeled CeSPD-2 ASH domain shows the conserved Immunoglobulin-like structural fold found in the overlapping homologous superfamilies of PapD, MSP and Chaperone Domains

The top ranked 3D model for the CeSPD-2 ASH domain was produced by LOMETS^42^ using the C-terminal PapD-like domain of human Hydin as a template (Figure 4A). The superimposed structures have overall high structural similarity (RMSD: 1.125 Å). Furthermore, the most conserved residues (F5, N22, and P53) have similar location and orientation in the superimposed CeSPD-2 and OCRL-1 ASH domains (Figure 4B). Although, the topology of Hydin and CeSPD-2 β-sheets differ, both domains form an antiparallel *β*-strands sandwich conformation (Figure 4C). Analysis predicted the formation of a classic type parallel *β*-bulge in both the CeSPD-2 ASH and the Hydin PapD-like domain (Figure 4C). The *β*-bulge in CeSPD-2 forms between residues A4, F5 on *β*1 and W91 on *β*7. To compare the CeSPD-2 ASH domain structure with other similar centrosomal proteins, the ASH domains of ASPM and ASP were also modeled (Figure 5). The structure of the ASH domain of OCRL-1 when bound to Rab8 (3QBT)^43^ was selected as the template for ASP, while the ASH domain of ASPM was modeled using the C-terminal domain of a PapD-like domain from an uncharacterized protein in *Porphyromonas gingivalis* (2QSV)^44^. A DALI structure-based search for structurally similar domains to the CeSPD-2 ASH domain fold identified ten domains of known structure (Figure 5B and Table 1). Like the sequence alignment of ASH domains in Figure 3A, the ten domains show little to no conservation of primary sequence amongst each other or with CeSPD-2 ASH domain. Despite this, all ten structures superimposed with low RMSD indicating a common structural fold (Figure 5B). The C-terminal PapD-like domain of HsHydin protein (2E6J^45^; Z-score: 15.4) and an ASH domain of HsCEP192 protein (6FVI^46^; Z-score: 11.1), were the two closest neighbors for the CeSPD-2 ASH domain (Figure 5B). Superimposition of the modeled CeSPD-2 ASH domain and the ten closest structural domain neighbors, shown in Movie 1, provides visual confirmation of how closely related the structure of these domains are.

**Figure 4.**
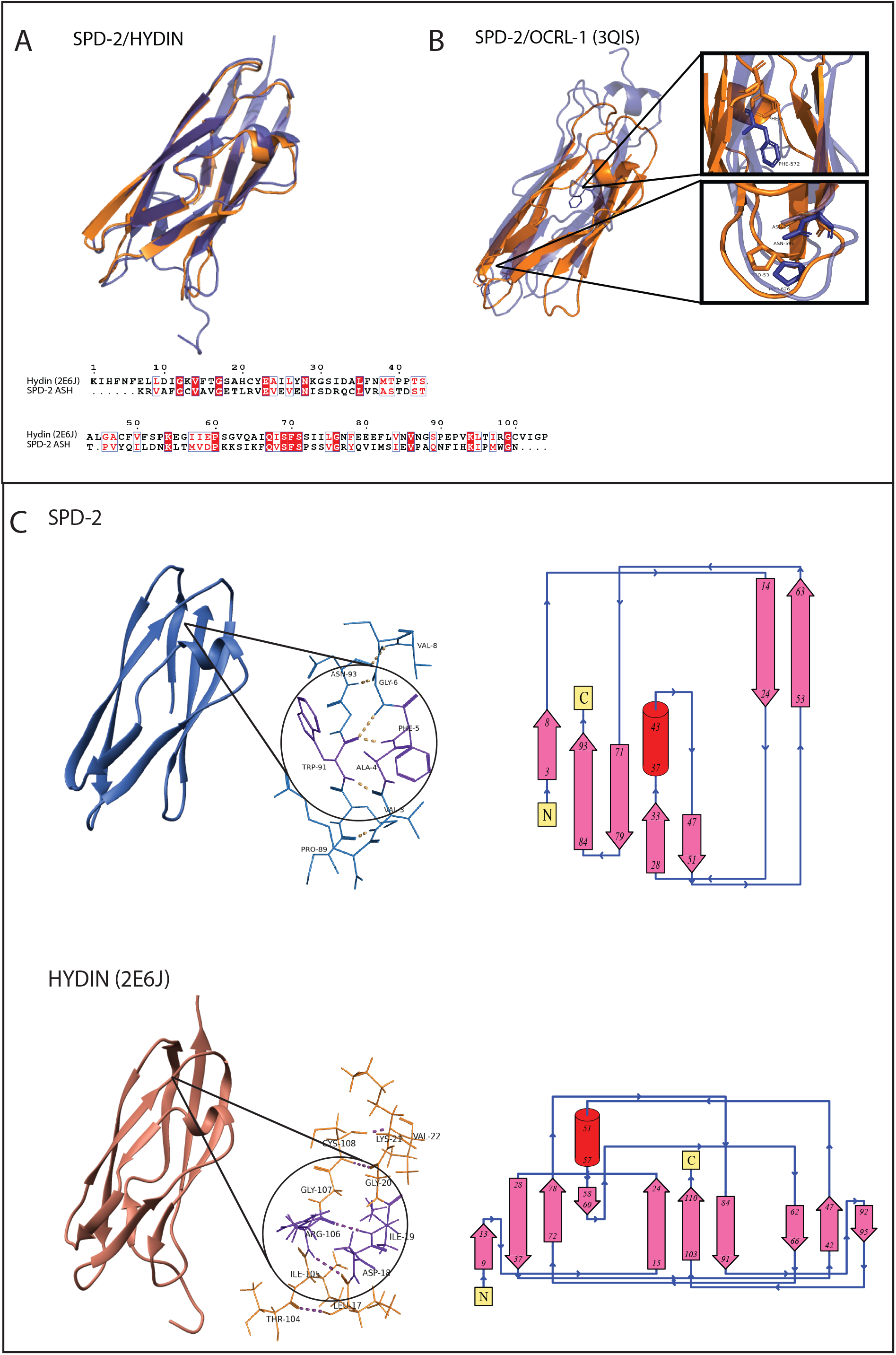
Comparison between the three-dimensional model of the SPD-2 ASH domain and the known structure of ASH domains of HYDIN or OCRL-1. A) Superposition of the modeled CeSPD-2 ASH domain with the human Hydin C-terminal PapD-like domain template (2E6J^45^) and sequence alignment based on the superposition. Residues with white letters that are highlighted in red are identical in both domains. Red letters in unfilled blue boxes correspond to similar residues. B) Overlays of SPD-2 ASH (orange) with the unbound structure of the ASH domain from OCRL-1 (3QIS^41^) with highly conserved residues highlighted and labeled. Images were produced with Pymol^137^. C) The three-dimensional ribbon diagram of the C-terminal Hydin PapD domain and the predicted CeSPD-2 ASH domains are shown next to their topology diagrams. The β-bulges in SPD-2 and Hydin are magnified, and participating residues are highlighted in purple. The arrows in the topology diagram represent β-sheets and their orientation. Cylinders represent α-helices, and the yellow boxes denote the N- and C-terminus of the domains. Residue numbers are included at the N- and C-terminus of each secondary structure.

**Figure 5.**
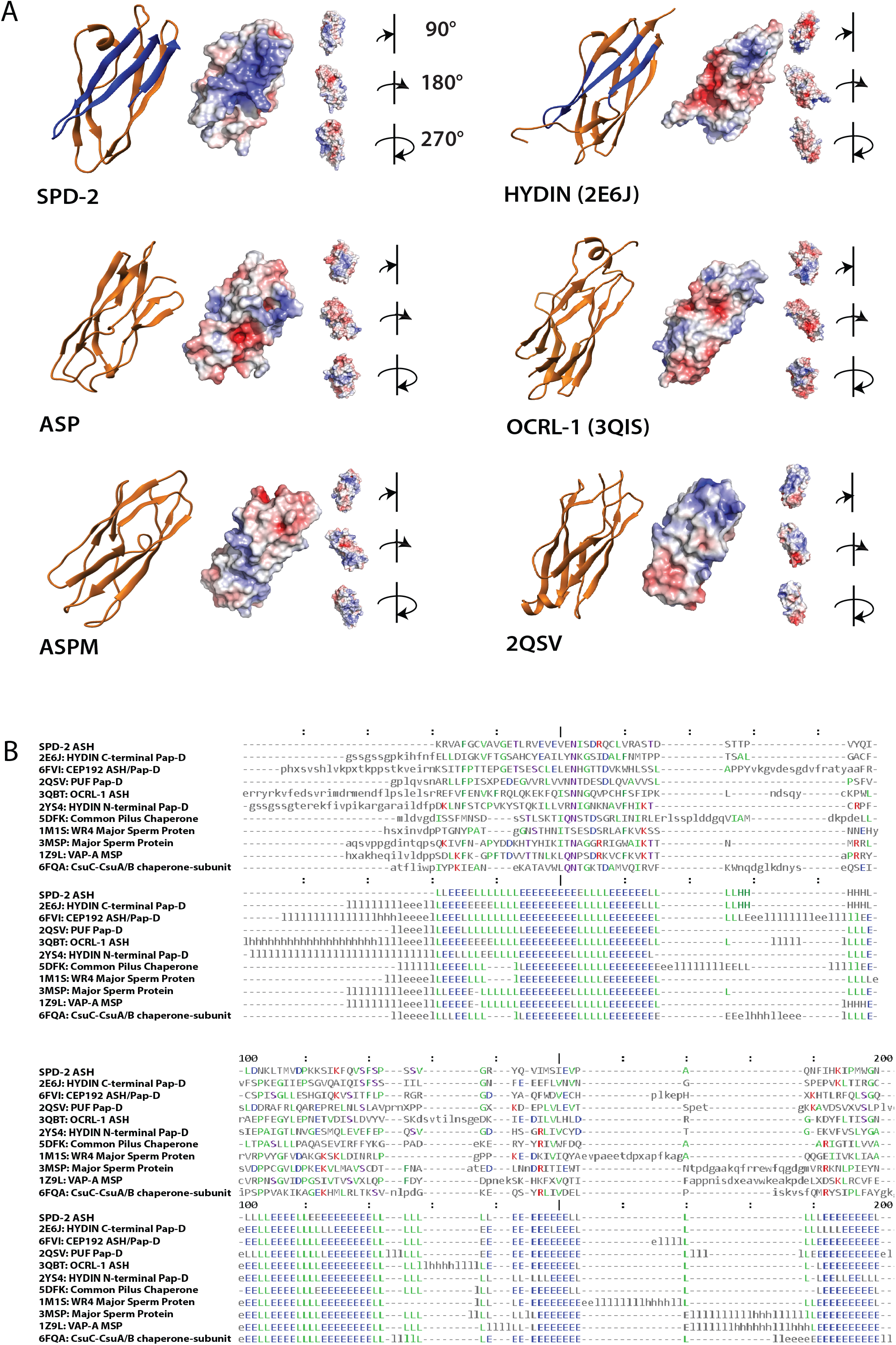
Electrostatic and structural similarities between the ASH domains from SPD-2 and other ASH domain containing proteins. The modeled structure of the SPD-2 ASH domain, the Human Hydin PapD-like domain (2E6J^45^), the Human OCRL-1 ASH domain (3QBT^43^), the Drosophila ASP ASH domain, and Human ASPM ASH domain are shown next to their surface electrostatics profiles mapped on to the models. The basic residues in CeSPD-2 and Hydin are highlighted in blue on the ribbon model. Structures for ASP and ASPM were modeled using OCRL-1 ASH when bound to Rab8A (3QBT^43^) and a protein of unknown function from *P. gingivalis* W83 (2QSV^44^) as templates, respectively. Three additional views of the electrostatic potential rotated clockwise by 90°, 180° and 270° are shown. Electrostatic images were produced with APBS-PDB2PQR server and PyMOL^137^, where blue represents basicity and red represents acidity (+5 to −5 kt/e). B) Structure based sequence alignment of structurally similar domains to CeSPD-2 identified using DALI^141^. The primary sequence and secondary structure of the domains similar to CeSPD-2 are shown in an alignment. The residues in the primary sequence in blue are the most conserved, residues in red show the lowest conservation and the residues in green are those with intermediate conservation. The secondary structure is annotated with β-sheets as blue letter E’s and the loops between them in green letter L’s.

### CeSPD-2 ASH Has a Unique Biochemical Character That May Have Functional Significance

To identify possible functional regions within the ASH superfold, we generated surface electrostatic potential maps for the predicted ASH domains of the centrosomal proteins CeSPD-2, ASP, ASPM and the domains that served as their structural templates (HYDIN, OCRL-1 and Porphyromonas gingivalis W83 protein of unknown function) (Figure 5A). The electrostatic profile of CeSPD-2 ASH shows a large basic groove that is formed by residues K1-C7 and Y71-N93 (Figure 5A). A much smaller basic groove is found at a similar location in the Hydin PapD-like domain (2E6J) and this groove is absent or significantly reduced in ASP, OCRL-1, ASPM, and 2QSV (Figure 5A). To assess functional significance of the basic groove on the CeSPD-2 ASH domain, we searched for putative protein-protein and protein-RNA interaction sites (Figure 6). Consistent with the possibility that the basic groove is functionally relevant, most of the residues in the basic groove (Figure 6A) are predicted to be involved in protein-protein interactions (Figure 6B). The predicted protein-RNA interactions (Figure 6C) also partially coincide with the basic surface (Figure 6A). Contact site predictions in other ASH domains are distinct from those in CeSPD-2 ASH (Figure S1). Given its proximity to and structural relatedness to the ASH domain, we generated a 3D model of the Ig-like domain in tandem with the ASH domain. The combined 3D model of the ASH and Ig-like domain shows that the basic region stretches from the cleft between the two domains to the outward facing surface of the first *β*-sheet (S2-K32), the cavity between the second *β*-sheet (K57-V63) and the short α-helix (M37-Q39) on the Ig-like domain.

**Figure 6.**
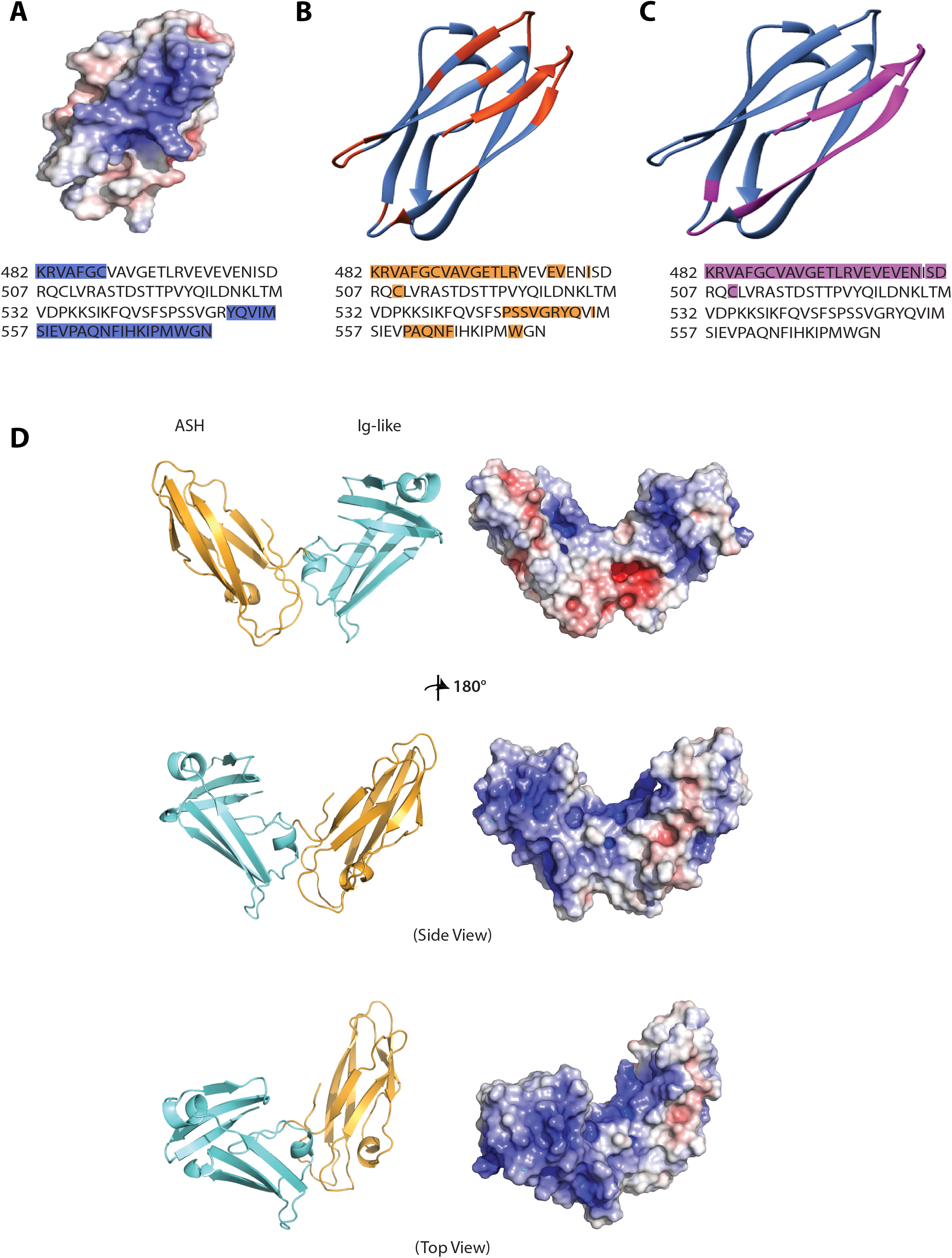
Predicted contact sites for SPD-2 ASH domain. A) The surface electrostatic profile of SPD-2 ASH with the corresponding residues that form the basic grove labeled in blue on the sequence below. B) Predicted structure-based protein-protein interaction sites are highlighted in orange on both the backbone of the modeled CeSPD-2 ASH domain and its sequence. C) Predicted RNA binding sites on the SPD-2 ASH domain are highlighted in purple in both the backbone structure and sequence. D) Model and electrostatic profile of the tandem ASH and Ig-like domains. The three-dimensional ribbon structure and predicted electrostatic profile of the SPD-2 ASH (gold) and Ig-like (blue) domain models assembled into a single unit. Images were produced with PyMOL^137^ and the APBS-PDB2PQR server. Blue represents basicity and red represents acidity (+5 to −5 kt/e).

### The CeSPD-2 ASH Domain Localizes to Centrosomes

Given that ASH domains in centrosomal proteins localize to the centrosome *in vitro*^47–49^, we hypothesized that ASH domain of CeSPD-2 localizes to the centrosome. To test this, we expressed the ASH domain of CeSPD-2 as a fusion protein with GFP *in vivo* (Figure 7). It should be noted that strains expressing ASH::GFP in the germline were hard to obtain and that germline transmission of the extrachromosomal arrays in these strains was very low (equal or less than 10%). This may suggest ASH::GFP construct may be toxic when overexpressed. When the construct was expressed in the germline (*pie-1P*::ASH::GFP), GFP was detected on the centrosomes and centrosome associated microtubules in cells of embryo undergoing mitosis (Figure 7F-H). Given the low transmission, low expression levels and the possible toxicity of the ASH::GFP expression in the germline, we also generated strains that conditionally expressed ASH::GFP upon heat-shock. Heat shock induced overexpression of ASH::GFP resulted in early embryos having a single GFP focus adjacent to the nucleus in interphase cells (Figure 7A-C) or several GFP aggregates of variable size within single cell (Figure 7D-E).

**Figure 7.**
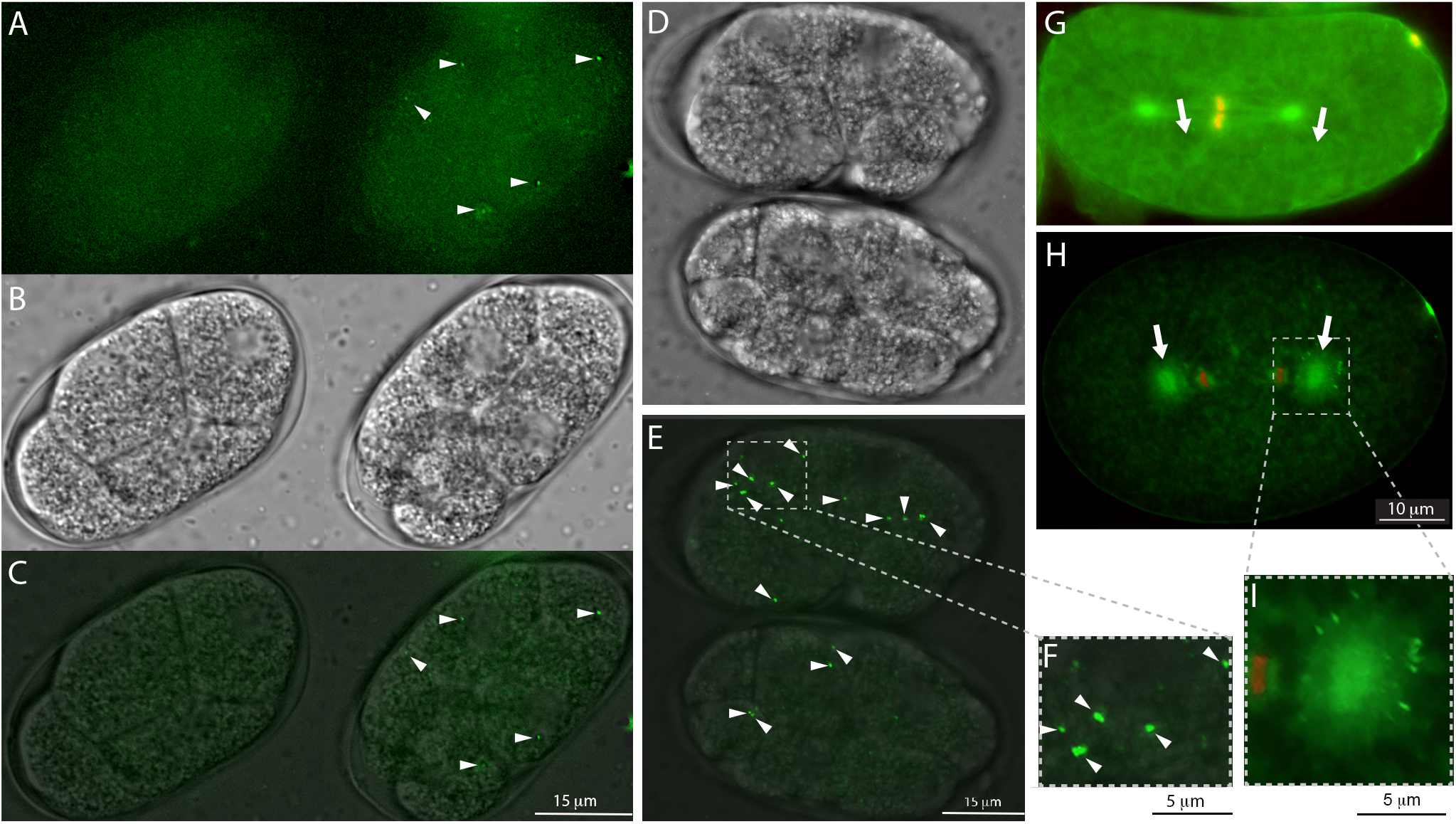
Overexpressing the SPD-2 ASH domain in *C. elegans* embryos. A-E) Images generated from partial z-stack projections of embryos expressing the translational fusion of the ASH domain and GFP under the control of an inducible heat-shock promoter. ASH::GFP is expressed from an extrachromosomal array that is inherited by a proportion of the embryos. Arrowheads mark ASH::GFP foci. Scale bar is 15 μm. A-C) Fluorescent (A), Differential Interference Contrast (DIC) (B) and merged (C) images of two embryos submitted to the same heat shock protocol. The embryo on the right expresses GFP that localizes in foci associated with cellular nuclei. Embryo on the left does not express ASH::GFP, likely because it did not inherit the extrachromosomal array including the ASH::GFP construct. D-F) A pair of embryos expressing ASH::GFP imaged by DIC (D) and fluorescence microscopy (E) merged fluorescence and DIC images. These embryos express higher levels of ASH::GFP that can be seen as larger foci. F) Digital magnification of a region of the embryo in E to highlight GFP aggregates within a cell. G-I) Images of embryos expressing MCherry::H2B (red fluorescent Histone marker) and ASH::GFP driven by the germline promoter P*pie-1*. ASH::GFP localizes broadly to the centrosome, spindle microtubules (G), and centrosome associated microtubule asters (H). Scale bar is 10 μm. Arrows point at the microtubule organizing center (centrosomes) in the dividing one cell embryo. I) is a digital magnification of a region of the embryo in I to highlight ASH::GFP localization on the centrosome and aster microtubules. Scale bar is 5 μm.

## Discussion

CeSPD-2 is a bi-functional protein that is essential for the non-overlapping roles regulating centriole duplication and centrosome maturation. The domain architecture predicted for CeSPD-2 is consistent with its known functions.

### CeSPD-2 intrinsic disorder may be key for its multifunctionality

The presence of a large, disordered N-terminal region followed by multiple structured domains in CeSPD-2 may provide mechanistic insight on its ability to act as a central regulator of multiple aspects of cell division. Interestingly, many intrinsically disordered proteins display phase transition behaviors^50^ and the PCM forms a compartment that appears to display such phase-separation properties^51^. In PCM reconstitution experiments SPD-5 forms spherical condensates that can be either initiated or enhanced by the SPD-2 and PLK-1 proteins^15^ and strengthen the SPD-5 condensates from mechanical disruption^52^. This may suggest the interesting possibility that the intrinsic disordered nature of SPD-2 may contribute to the liquid-liquid phase-separated nature of the mitotic PCM. Alternatively or additionally, SPD-2 intrinsic disordered regions may provide the conformational plasticity to act as a hub for protein interactions similarly to many important regulators of transcription, translation and cell cycle that have intrinsically disordered regions^53,54^. Protein modifications including phosphorylation within disordered regions often change the signaling output of proteins and can function as switches at a specific threshold of phosphorylation^53,55^. We speculate that the disordered regions in CeSPD-2 may be key for its ability to accommodate multiple binding partners, operate in different processes, and mediate the link between the process of centriole duplication and centrosome maturation. Most CDK targets contain multiple phosphorylation sites within intrinsically disorder regions^56^ that allow for multi-phosphorylated outputs^57^. Specific signal outputs of Cdk1 targets can be extended to other kinases that use phosphorylated CDK sites as priming sites as is the case with PLKs^57,58^. This has been proposed to be a general mechanism for CDK producing specific ultrasensitive responses in its targets^55,57^. The N-terminal disordered region of CeSPD-2 contains several predicted and experimentally confirmed CDK and PLK phosphorylation sites ^3,10,59^. A tantalizing possibility is that phosphorylation of CeSPD-2 at specific CDK sites may result in conformational changes that promote its binding to either PLK-1 or PLK4/ZYG-1 during centrosome maturation and duplication respectively. CDK is a limiting factor for centrosome growth during mitosis as it helps initiate mitotic PCM assembly^10,60,61^, and is also part of a kinase hierarchy that regulates centrosome duplication^2,62–65^. Specific CDK phosphorylation may prime PLK-1 binding to the N-terminus of CeSPD-2^59^, which is required for recruitment to the centrosome and phosphorylation of SPD-5^10^. A different CDK phosphorylation signature may prime SPD-2 N-terminus for PLK4/ZYG-1 binding^66^, an interaction required for the initiation of centriole duplication^28,67,68^.

### The ASH Domain targets SPD-2 to the centrosome

Our finding that the CeSPD-2 ASH domain alone is sufficient for localization to the centrosome *in vivo* is consistent with prior observations that mutations in ASH domains impair SPD-2/CEP192 localization to the centrosome and cilia^2,48,69^, and supports similar observations *in vitro*^34,47–49,70^. Interestingly, localization of the ASH domain in embryos differs depending on the levels of its expression (Figure 7). Expression of ASH driven by the *pie-1* promoter throughout germline development results in ASH::GFP localization to the centrosome and centrosome-associated microtubules during mitosis. However, rapid overexpression of ASH protein in animals carrying a heat shock inducible ASH::GFP construct results in a GFP focus associated with interphase nuclei and larger in size cytosolic GFP aggregates. Interestingly, a phospho-mimicking mutation S545E within the ASH domain results in supernumerary centrioles in intestinal cells^69^. This variable pattern of ASH::GFP localization may reflect activity or stability altering modification(s) and/or thresholds at which the ASH domain may have different functions.

### The “SPD-2 Domain”: ASH and Ig-like Domain Function

The *C. elegans* SPD-2 ASH domain is located within a region that has been termed the “SPD-2 domain” extending from K482 to R714 that also includes the Ig-like domain and the beginning of the PDZ-like^3^. A point mutation on the Ig-like domain (G615) results in the lack of a pseudocleavage furrow, incorrect localization of the sperm pronuclei, and bipolar spindle assembly defects^2^. These defects suggest that the Ig-like domain may play a functional role in PCM maturation and microtubule nucleation. Our 3D structure prediction of the ASH domain and neighboring Ig-like domain together (Figure 6D), resemble the structure of bacterial PapD-like chaperones that facilitate the assembly of both pilus and non-pilus organelles^71,72^. These chaperones consist of two Ig-like domains oriented toward each other in an L-shape and the cleft region between the two domains form a pilus subunit binding pocket^73^. Based on the mechanism of function of similarly structured intramolecular chaperones^74^, we propose that the Ig-like and ASH domains may regulate exposure of the reactive basic pocket formed in the cleft between these domains. An interesting possibility is that this potential chaperone mechanism could be regulating ZYG-1 recruitment to the centriole by controlling access to highly charged ZYG-1 interacting residues located at the N-terminus of CeSPD-2. Shimanovskaya and colleagues (2014) ^67^ found that changes to this region on CeSPD-2 disrupted ZYG-1 docking and centriole duplication, but not CeSPD-2 localization. The authors proposed that availability of the acidic region in CeSPD-2 may be regulated to ensure that ZYG-1 interacted specifically with centriolar CeSPD-2 and not centrosomal CeSPD-2.

### Coiled-coils and GEF domain’s possible role in zygote polarity establishment or maintenance

CeSPD-2 coiled-coil has been reported to have three coiled-coil domains^2^. Our analysis revealed four coiled-coils. The first two coiled-coils (CC1 and CC2) overlap with those previously reported, however in our prediction the third coiled-coil (CC3) is slightly shifted towards the C-terminus. The predicted coiled-coils CC3 and CC4 are located within a predicted GTP exchange factor (GEF) motif (Figure 2). This GEF signature is structurally most similar to an unconventional group of GEFs typified by the *S. cerevisiae* Sec2p GEF motif that contains large coiled-coils responsible for activation of the Sec4p Rab GTPase that regulates polarized exocytosis in yeast^75,76^. An exciting possibility is that this GEF domain could be involved in establishing and/or maintaining zygote polarity that determines the anterior-posterior axis of the one-cell embryo in *C. elegans*^8,77^. Several *spd-2* mutants have polarity defects including abnormal distribution of cell cortex enriched PAR (abnormal embryonic PARtitioning of the cytosol) proteins.^8^ PAR proteins are evolutionarily conserved regulators of cell polarity and polarized vesicle trafficking^78,79^ and rely on small GTPases for their dynamic localization^80–83^. The initial asymmetry in cortical PAR protein distribution in the zygote is triggered soon after fertilization^8,84^, and relies on an anterior to posterior cytoplasmic flow achieved by asymmetric contraction of the actomyosin cytoskeleton that is inhibited in the proximity of the paternal pronucleus by the sperm derived mature centrosome pair^83^. One of the cues provided by the centrosome is AIR-1^85,86^, the Aurora A kinase recruited by SPD-2 to the paternally provided centrosome. AIR-1/Aurora A kinase, signals posterior fates in the cortex closest to the centrosome associated sperm pronucleus in a microtubule independent manner^80,86,87^ by preventing the RhoGEF ECT-2 from localizing to the zygote’s posterior cortex in the vicinity of the sperm pronucleus associated centrosome, and thus limiting RHO-1 function to the anterior of the embryo. ECT-2 affects the localization of PAR proteins by affecting actomyosin network dependent cortical contractibility^80,87^. The presence of the putative GEF domain in CeSPD-2 raises the possibility that CeSPD-2 may have a role in zygotic anteroposterior patterning independent of its role recruiting AIR-1 to the centrosome. The identification of *spd-2* in unbiased RNAi screens for polarity proteins involved in endocytic traffic^79^ supports the possibility that CeSPD-2 could potentially regulate a Rab GTPase(s) given its similarity of its GEF domain with the *S. cerevisiae* Sec2p, which regulates Rab GTPase polarized vesicle trafficking. In *Drosophila*, localization of Bazooka/PAR-3 to the anterior cortex of the oocytes occurs in two phases that involve vesicular trafficking. The first phase requires the small GTPases RAB5 and the second phase requires RAB11, microtubules, the dynein motor, and IKK-related kinase (IKKε)^88^. Interestingly, in *C. elegans*, *rab-5 and rab-11*mutants have cell polarity phenotypes including mislocalization of cortical PAR proteins, and pronuclear migration and rotation^82,87,89–91^. It is also possible that the CeSPD-2 GEF domain is inactive but still binds and regulates a Rab GTPase, as is the case with the GAP domain in the ASH containing protein OCRL-1^92,93^.

### Role of the PDZ-like domain: protein-protein and intramolecular interactions

At the C-terminus of CeSPD-2, we identified a previously unreported putative PDZ-like domain. PDZ domains are one of the most common protein-protein interaction modules involved in cellular signaling. These domains are involved in both intra- molecular and inter-molecular interactions that provide functional diversity^33^. A point mutation in CeSPD-2 PDZ domain, results in embryos that lack centrosomes, exhibit bipolar spindle defects and polarity defects that result in incorrect sperm pronucleus migration and failure of pronuclei rotation, as well as abnormal localization of PAR proteins^2,8,69^. These phenotypes may indicate that the PDZ-like domain interacts with alternative partners for regulating centrosome maturation and embryo polarity. One interesting possibility is that, like PDZ-RhoGEF proteins, SPD-2’s PDZ-like domain may impact GTPase mediated microtubule dynamics^94^ and actin cytoskeletal reorganization^95^ thereby impacting interactions with the dynein heavy chain *dhc-1* motor required for normal levels of CeSPD-2 localization to the centrosome, centrosome separation, pronuclear migration, vesicle transport, and centrosome cortex interactions during polarity polarity establishment in the one cell embryo polarity^2,96,97^. It is also possible that the PDZ-like domain might interact with other domains within CeSPD-2. The polarity defects in animals with mutations in the PDZ-like domain^2^ may support possibility of intra-molecular interactions between this domain and the GEF domain.

### Identification of the ASH/PapD Superfold

Despite low sequence similarity (≤ 34%) and differing topologies, ASH domains share a highly conserved immunoglobulin-like fold that appear to be most structurally related to the first domain of PapD/PapD-like chaperones. We were unable to find unique structural features in ASH domains that clearly differentiate them from PapD/PapD-like domains. This is likely why ASH and PapD/PapD-like domains are often used interchangeably. Like ASH and PapD/PapD-like domains, MSP domains also belong in the conserved family of immunoglobulin-like structural fold despite only sharing 11% sequence similarity with PapD/PapD-like folds^98^. The similarity between nematode MSP domains to PapD chaperone domains^71,99^ is thought to be due to the potential of MSPs to bind interfaces used for filament biogenesis and alter the rate of filament formation^100^. Although not explored from a functional standpoint in previous studies, it is clear that all of these domains belong to the same superfold within the larger Immunoglobulin superfamily. MSP domains are classified as members of the PapD-like superfamily, which overlaps with the Ig-like superfamily. Therefore, ASH domains generally fall under the Ig-like fold category, but there is currently not enough data to define the ASH domains as its own structural family.

## Conclusion

Here we identify the SPD-2 ASH domain as a potential module designed to target proteins that interact with SPD-2 to the centrosome, and as an ideal small peptides tool to query mechanisms of centrosome inheritance and function, previously lacking in *C. elegans*. We speculate and propose possible mechanisms for predicted domains in SPD-2 and suggest putative domains for uncharacterized protein interactions as well as for known SPD-2 interacting partners. This study lays the groundwork for designing rational hypothesis-based experiments for future analyses that probe the mechanisms of SPD-2 function *in vivo*.

## Materials & Methods

### Secondary Structure Prediction and Sequence Analysis

The available full-length *C. elegans* SPD-2 protein sequence was retrieved from the NCBI Reference Sequence (RefSeq) Protein sequence database (NP_492414.1; The *C. elegans* Sequencing Consortium, 1998^101^). The domain architecture for SPD-2 was analyzed using SMART (Simple Modular Architecture Research Tool) ^102^, HHpred (v.2016) ^103^, Pfam^104^, Prosite^105^, Interpro^106^, and the NCBI Conserved Domain (CD) database^107^}, and a final consensus domain architecture cartoon was drawn to scale using Adobe Illustrator (Adobe Inc., 2019). Predictive secondary structure analysis of the full-length protein and ASH domain specific sequence was used to confirm domain boundaries and assess the underlying secondary structure using PSIPRED^108^, PSSpred^109^, SPIDER2^110^, Jpred4^111^, and SABLE^112^. Based on the domain architecture and secondary structure prediction results, the following sequence analysis was also performed for a more detailed sequence characterization: (a) Sequence disorder was predicted using Genesilico MetaDisorder^113^ and PrDOS^114^, and (b) Coiled-coils were predicted using COILS^115^, DeepCoil^116^, Paircoil2^117^, and Multicoil^118^. To assess potential sequence-based intermolecular interaction sites, ANCHOR^119^, DisoRDPbind^120^ were used. DisoRDPbind predicts interaction sites within intrinsically disordered regions.

The ASH domain protein sequences of additional related proteins were retrieved from NCBI: Human [Hs] hydrocephalus-inducing protein/HYDIN isoforms a/b and c/d (NP_001257903.1:513-613, NP_001185471.1:541-640), abnormal spindle-like microcephaly-associated protein/ASPM isoform 1/2 (NP_. 060606.3:37-134), inositol polyphosphate 5-phosphatase/OCRL-1 isoform a (NP_000267.2:564-678) and centrosomal protein 192kDa/CEP192 (NP_115518.3:2256-2402); *Xenopus laevis* [Xe] centrosomal protein/CEP192 (ADP36856.1:1776-1846); Mus musculus [Mm] hydrocephalus inducing protein/HYDIN (NP_766504.3:559-659); *Drosophila melanogaster* [Dm] abnormal spindle protein (NP_524488.3:25-122). Domain sequence alignments were made using the MUSCLE webserver^121^ and annotated with ESPript3^122^. A sequence similarity matrix was constructed using the SIAS (Sequence Identity and Similarity) webserver^123^.

### Modeling and Structure-based Analysis

Human Hydin PapD like C-terminal ASH domain (2E6J^45^) was the best candidate template identified for modeling the SPD-2 ASH domain using various fold recognition algorithms. Three dimensional structural models of the CeSPD-2 ASH domain were generated using template-based modeling programs as well as *ab initio* approaches with Modeller^124^, RAPTOR-X^125^, Phyre2^126^, I-TASSER^127^, LOMETS^42^, and Swissmodel^128^. Model quality was evaluated using ProSA-Web^129^, Verfiy3D^130^, ProQ3^131^, and VoroMQA^132^. The model with the best evaluation profile was further refined using ModRefiner^133^ and used for all subsequent analyses. The *ab initio* modeling algorithm in Robetta^134^ was also used to gather predicted structural data for the disordered domains in SPD-2. DEMO^135^ was used to assemble full-length models. Electrostatic potential maps were produced and visualized using the Adaptive Poisson-Boltzmann Solver (APBS) - PDB2PQR server^136^ and the PyMOL (v.1.8) molecular graphics system^137^. Chimera (v1.11)^138^ was used for structure alignment and the production of backbone images. Potential contact sites based on the sequence and predicted structure of the model was predicted using SPPIDER (Solvent accessibility-based Protein-Protein Interface iDEntification and Recognition)^139^. This procedure was repeated for modeling the ASH domains of *Drosophila* abnormal spindle protein/ASP and human abnormal spindle-like microcephaly-associated protein/ASPM.

### Plasmid Construct

The *pie-1P*::ASH::wGFP clone was generated by synthesis (Genewiz).This construct includes the *pie-1* gene promoter in the pCM1.127 plasmid (Addgene), driving a translational fusion of the SPD-2 ASH domain (nucleotides 1444-1720) and the wGFP sequence isolated from the pCFJ1848 plasmid (gift from Erik Jorgensen). The 3’UTR in this construct contains the *pie-1* 3’UTR from the pCM5.47 plasmid (Addgene). The *hsp-16.2*P::ASH::GFP construct was generated by cloning the SPD-2 ASH domain sequence (nucleotides 1444-1720) into the PTB11 vector (Andrew Fire plasmid kit). This vector includes the hsp-16.3 promoter and the unc-54 3’UTR.

### Culture Conditions and Strains

Worms were cultured using standard techniques^140^. All strains were maintained at 16°C unless otherwise noted. The *pie-1*p::ASH::GFP plasmid was injected (15ng/uL) into the OD95 strain - unc-119(ed3) III; Itls37 IV[(pAA64) pie-1p::mCherry::his-58 + unc-119(+)]; ltIs38 III [pie-1p::GFP::PH(PLC1delta1) + unc-119(+)] by NemaMetrix (Murray, UT). A *rol-6* construct (pNU406) was used as the injection marker. The MSC25 strain that resulted from these injections transmits the extrachromosomal to about 10% of its progeny. MSC25 roller hermaphrodites were crossed with AV675 [mCherry::H2B] males and the resulting progeny was screened for ASH::GFP expression. The *hsp-16.2*p::ASH::GFP plasmid (2-10ng/ml) was co-injected with the *rol-6* (PRF4) injection marker (15-25ng/uL) into the germline of N2 nematodes. The resulting strain (MSC26), expressed ASH::GFP upon heat shock treatment.

### Heat Shock Treatment, Injection and Gonad Dissection

Gravid MSC6 hermaphrodites were heat-shocked at 30 °C for 4 hours. After heat-shock, nematodes were recovered at 16°C for 1-2 hours. After recovery gravid MSC26 hermaphrodites were placed in 2-5 μl drop of M9 on a glass slide cover slip and were dissected to release the embryos. Coverslips with dissected MSC25 or MSC26 hermaphrodites were placed onto microscope slides with a 4% agar cushion sealed with Vaseline, and embryos were immediately imaged.

### Imaging

Z-stacks images of embryos were collected with the DeltaVision Deconvolution system mounted on an Olympus IX-70 inverted microscope equipped with a CoolSnap HQ CCD camera and a LED arc bulb illumination source, using 60×1.514N oil immersion objective, GFP/mCherry fluorescence filter sets, and DIC optics. Image processing and z-stack projections was done with the SoftWorx software included in the Delta Vision Deconvolution system.

## Abbreviations

AIR: Aurora kinase
ASH: Aspm-SPD-2 Hydin
ASP: Abnormal Spindle Protein
ASPM: Abnormal Spindle-like Microcephaly-associated Protein
CC: coiled-coil
CDK: Cyclin dependent Kinase
Ce: Caenorhabditis elegans
CEP: Centrosomal Protein
CPAP: centrosomal P4.1-associated protein
D: Drosophila
GAP: GTPase activating protein
GEF: GTPase guanine nucleotide exchange factor
Hs: Homo sapiens/Human
Ig: Immunoglobulin
MAP: Microtubule associated Protein
MSP: Major Sperm Protein
MDP: Major Sperm Domain-Containing Protein
OCRL-1: Golgi endocytic trafficking protein Inositol polyphosphate 5-phosphatase
PAR: abnormal embryonic PARtitioning of the cytosol
PCM: Pericentriolar material
PCMD: pericentriolar matrix deficient
PDZ: PSD95/Dlg-1/zo-1
PLK: Polo like kinase
RMSD: Root Mean Square Deviation
SAS: <u>S</u>pindle <u>as</u>sembly abnormal proteins
SPD: Spindle-defective protein
TRAPP: TRAnsport Protein Particle
Xe: Xenopus
ZYG: zygote defective protein

## Acknowledgements

We thank the Caenorhabditis Genetics Center funded by National Institute of Health (NIH) Office of Research Infrastructure Programs P40OD010440) for strains. The authors would like to thank Tamara Mikeladze-Dvali and Elif Nur Firat Karalar, for providing us with constructive feedback. This work was supported by a PSC-CUNY award TRADB-46-113 and NIH grant 1SC2GM118275-01.

