## Supplemental table 1 for "The Centrosomal Swiss Army Knife: A combined in silico and in vivo approach to the structure-function annotation of SPD-2 provides mechanistic insight into its functional diversity"

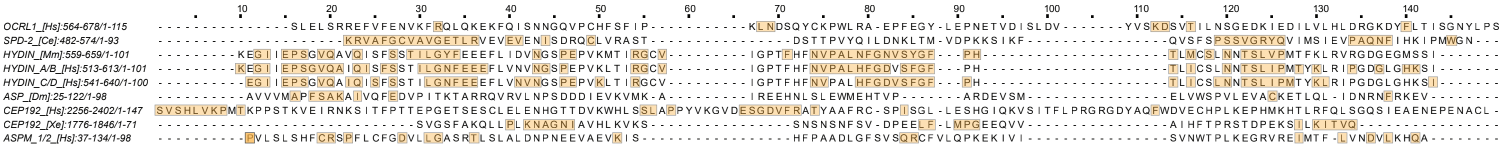


**Suppl. ASH Domain Contact Sites**

Multiple sequence alignment of ASH domains (see Figure 3A). The structure-based contact site predictions for each sequence are boxed and highlighted.
